# comradesOO: An Object-Oriented R Package for the Analysis of RNA Structural Data Generated by RNA crosslinking experiments

**DOI:** 10.1101/2023.12.12.563348

**Authors:** Jonathan L. Price, Omer Ziv, Malte L. Pinckert, Eric A. Miska

## Abstract

**Summary:** RNA (Ribonucleic Acid) molecules have secondary and tertiary structures *in vivo* which play a crucial role in cellular processes such as the regulation of gene expression, RNA processing, and localization. The ability to investigate these structures will enhance our understanding of their function and contribute to the diagnosis and treatment of diseases caused by RNA dysregulation. However, there are no mature pipelines or packages for processing and analysing complex *in vivo* RNA structural data. Here, we present COMRADES Object-Oriented (comradesOO), a novel software package for the comprehensive analysis of data derived from the COMRADES (Crosslinking of Matched RNA and Deep Sequencing) method. comradesOO offers a comprehensive pipeline from raw sequencing reads to the identification of RNA structural features. It includes read processing and alignment, clustering of duplexes, data exploration, folding and comparisons of RNA structures. comradesOO also enables comparisons between conditions, the identification of inter-RNA interactions, and the incorporation of reactivity data to improve structure prediction.

**Availability and Implementation:** comradesOO is freely available to non-commercial users and implemented in R, with the source code and documentation accessible at [https://CRAN.R-project.org/package=comradesOO]. The software is supported on Linux, macOS, and Windows platforms.

**Supplementary information:** Supplementary data are available at *Bioinformatics* online.

## 1 Introduction

RNA molecules exhibit secondary and tertiary structures *in vivo*. While ribosomal RNA (rRNA) with secondary structure and base pairings between nucleotides is a familiar concept, mRNA is frequently represented visually as a linear entity, typically marked with 5′ and 3′ labels (Vicens and Kieft, 2022). This bias in conceptualisation, compounded by the complexities of investigating RNA structures *in vivo* has led to the study of RNA structure lagging behind other fields of structural biology.

RNA structure is observed as dynamic *in vivo*, adapting to localized spatiotemporal conditions within the cell (Solayman *et al*., 2022). Factors such as minor changes in pH, salt concentrations, ligand availability, temperature or point mutations can influence the behavior of covalent base pairs, consequently affecting the structure (Wan *et al*., 2011). These structural changes have a diverse impact on cellular biology (Mortimer, Kidwell and Doudna, 2014), including transcriptional regulation (Tsai *et al*., 2010), splicing (Kar *et al*., 2011), translation (Ray *et al*., 2009) and RNA decay (Fukuchi and Tsuda, 2010).

Studying RNA structure *in vivo* is becoming a combinatorial assay with recent success in the field coming from utilising psoralen crosslinking methods, chemical probing and *in silico* folding for the same RNA (Spitale and Incarnato, 2023). This is because the limitation of each of the methods are mitigated by the others; psoralen crosslinking methods such as COMRADES (Crosslinking of Matched RNA and Deep Sequencing) (Ziv *et al*., 2018, 2020), PARIS (Lu, Gong and Zhang, 2018), SPLASH (Aw *et al*., 2016), LIGR-seq (Sharma *et al*., 2016), provide evidence for long-range base-pairing although not at base-pair resolution which complicates the use of in silico folding methods. Chemical probing methods, such as icSHAPE (Flynn *et al*., 2016), when applied alone, are limited by the presence of RNA binding proteins, solvent accessibility and their inability to detect long-range base pairing. However, their ability to provide information at single nucleotide resolution can improve in silico folding predictions of RNA crosslinking data.

The subsequent analysis of this combinatorial data is complicated, and no mature pipelines or packages exist. Furthermore, current analysis methods require multiple disjointed steps and difficult installation procedures. To this end, here we present the comradesOO R package (Crosslinking of Matched RNAs And Deep Sequencing - Object Oriented). comradesOO offers a complete pipeline, from raw reads to folded structures, although the package was designed to analyse COMRADES data specifically, the comradesOO R package will accept any psoralen crosslinking data presented in the correct format. The object-oriented nature of the package allows the storage of raw and processed data, as well as the use of methods which act on the object and perform many of the common procedures in this type of data-analysis.

## 2 Methods and Application

### 2.1 Read pre-processing

The COMRADES experimental protocol results in high-throughput sequencing data in FASTQ format. To process these raw sequencing reads and produce aligned reads for downstream analysis with the comradesOO package, we have developed a Nextflow (Di Tommaso *et al*., 2017) pre-processing pipeline Parameters for steps in the Nextflow pipeline can be found in **Supplemental Table 2** (https://github.com/JLP-BioInf/comradesNF). Crosslinking experiments have varied library preparation protocols and often small differences mean that it is not possible to follow a prescribed pipeline for data pre-processing. For this reason, users can also create their own input files provided they follow the guidelines set out in the vignette and **Supplemental Table 3**.

### 2.2 Overview of the comradesoo package

The comradesOO package can be installed using *install*.*packages* in R. A full vignette and usage documentation can be found on the CRAN website (https://CRAN.R-project.org/package=comradesOO) and through the *vignette* function. The object-oriented R package centers around a new S4 class the *comradesDataSet*. The class has slots that facilitate the storage and accessibility of COMRADES data, a more detailed explanation of the attributes can be found in the vignette (**Figure 1 a**).

**Figure 1.**
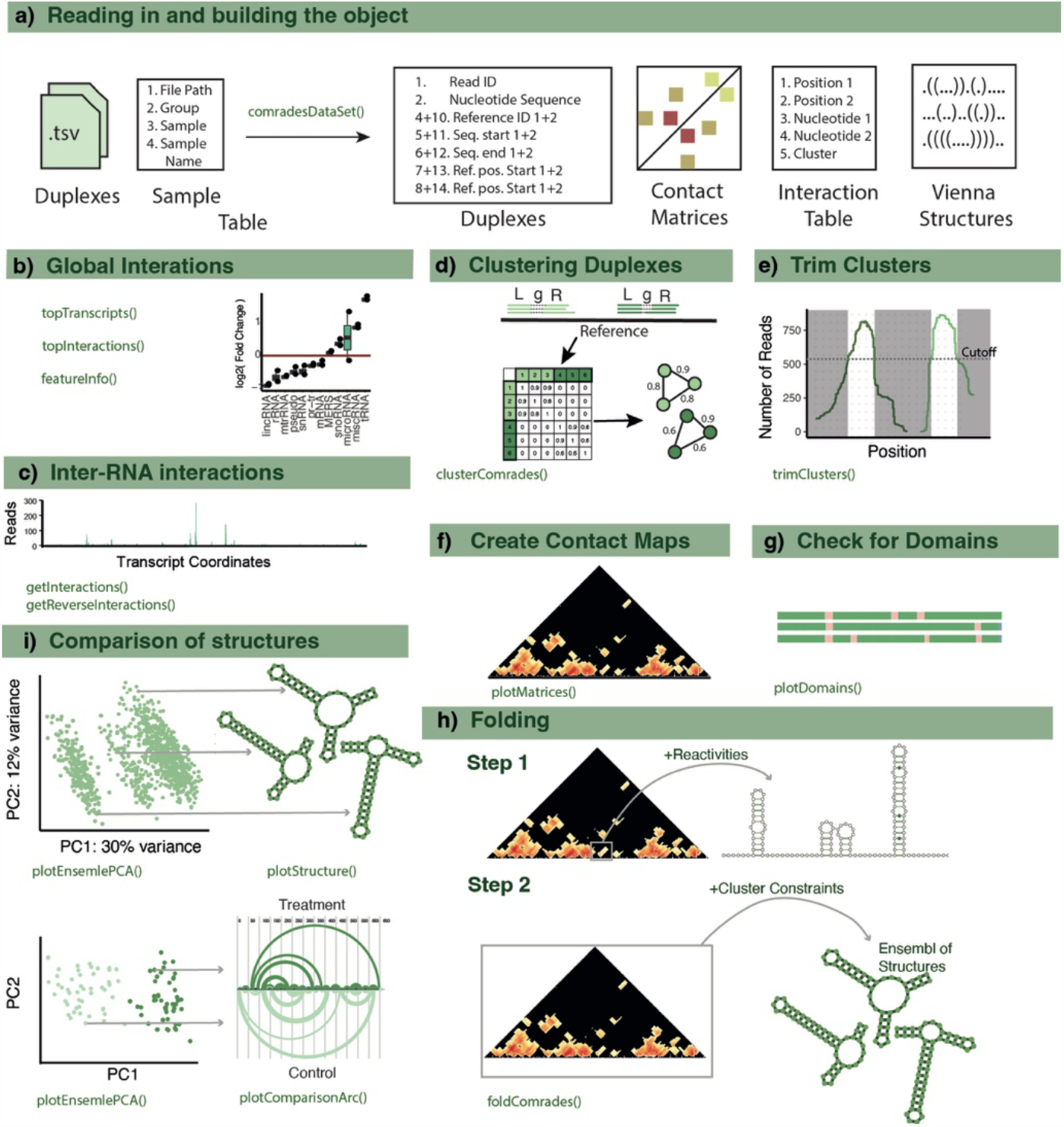
Schematic of comradesOO. **a)** Input for comradesOO and main slots of the comradesDataSet object. **c-j)** Schematics and examples of steps in the comradesOO package, further detail for each step can be found in the main text.

### 2.3 Reading in and exploring the global interactions

Loading the data into the comradesOO pipeline requires; 1) Sample metadata in the form of a tab-separated table. 2) The output of the COMRADES Nextflow pipeline or files in the same format (**Supplemental Table 3**). 3) The ID of the RNA of interest. The COMRADES experimental protocol involves a round of enrichment for a specific RNA. However, the resulting data also contains structural information from other RNAs, as well as inter- and intra-RNA interactions for the RNA of interest. To gain a comprehensive overview of the RNAs within the dataset, there are three primary methods: *featureInfo, topTranscripts* and *topInteractions*. These methods present the user with a table showing highly abundant RNAs and RNA-RNA interactions in the data set (**Figure 1 b**).

### 2.4 Exploring the inter-RNA interactions

The COMRADES protocol crosslinks any nucleotide bound RNAs, this includes inter-RNA interactions. Exploring these interactions after using the *topInteractions* and *topInteractors* methods can be performed with, *getInteractions* and *getReverseInteractions*. Plotting the resultant tables shows the location of reads for another chosen transcript (**Figure 1 c**).

### 2.5 Exploring the intra-RNA interactions

#### 2.5.1 Clustering and Trimming duplexes

In the COMRADES data, crosslinking and fragmentation leads to the production of redundant structural information, where the same *in vivo* structure from different RNA molecules produces slightly different RNA fragments. Clustering of these duplexes that originate from the same place in the reference transcript reduces computational time during the folding step and allows trimming of these clusters to improve the resolution. Clustering is performed as described in (Ziv *et al*., 2020). Briefly, gapped alignments can be described by the transcript coordinates of the left (L) and right (R) side of the reads and by the nucleotides between L and R (*g*). Reads with similar or identical *g* values are likely to originate from the same structure of different molecules. In comradesOO, an adjacency matrix is created for all chimeric reads based on the nucleotide difference between their g values. From these weights the network can be defined as: G = (V, E). To identify clusters within the graph the graph is clustered using random walks with the *cluster_waltrap* function (steps = 2) from the iGraph package (Csárdi *et al*., 2023). These clusters often contain a small number of longer L or R sequences due to the random fragmentation in the COMRADES protocol. Given the assumption that the reads within each cluster likely originate from the same structure in different molecules these clusters can be trimmed to contain the regions from L and R that have the most evidence (**Figure 1 d, e**) the clustering and trimming is achieved with the *clusterComrades* and *trimClusters* methods.

#### 2.5.2 Check for Domains

Folding RNAs in silico is computationally more expensive and inaccurate as the size of the RNA increases. To allow the user to fold smaller parts of the RNA of interest comradesOO has the *plotDomains* method. In the analysis of Hi-C data, domains are used to compartmentalise areas of the DNA with high inter-domain interactions and less interactions outside of the domain. Here we utilise a package designed for Hi-C analysis, TopDom (Shin *et al*., 2016) to achieve this effect (**Figure 1 g**).

#### 2.5.3 Folding

After choosing a domain the user can create predicted structures for any region or the whole of the RNA of interest using the *foldComrades* method. The folding works as follows; firstly, all clusters in the region are folded *in silico* using RNAFold from the Vienna package (Lorenz *et al*., 2011). For short range clusters (gap > 10 nt) this is done by folding the region with RNAFold. For long range clusters, an artificial linker is created between the two sides of the cluster and then this sequence is folded using RNAFold. If chemical probing data is available, the reactivities are included here. From these predicted structures of the clusters, the nucleotide contacts are then stored as constraints for the next step in the folding. Due to alternative topologies of the RNA, some of these predicted interactions maybe mutually exclusive. In step two the transcript region is folded a number of times to produce a representative structural ensemble. Each time the RNA is folded, hard constraints that were identified in the first step are added sequentially and the RNA is refolded. In the case where a constraint that is added shares a nucleotide position with a previously added constraint this new constraint is simply removed, and a new constraint is added. The user specifies how many constraints are added to each of the folded molecules. This produces an ensemble of structures that is stored in the object (**Figure 1 h**). To aid the analysis of the representative structural ensemble there are 3 functions; *plotEnsemblPCA, plotComparisonArc, structurePlot* (**Figure 1 i**).

### 2.6 Usage of comradesOO on ribosomal RNA dataset

To demonstrate the functionality of comradesOO, **Supp.Figure 1** shows the analysis of the 18S ribosomal rRNA. The inter-RNA interaction with the most evidence is the interaction between the 18S and 28S rRNA shown in **Supp. Figure 1 A**. Domain identification identifies several domains in the 18S RNA which can be taken through to the folding step, and these agree with the crystallography structure (Ban *et al*., 2000) (**Suppl. Figure 1 B, Supp. Table 2a**). Clustering of the data identified 83, 79 and 85 trimmed clusters for samples 1, 2 and 3 respectively with 78, 77, and 80 percent able to be explained by Watson and Crick base pairs in the crystallography structure **Supp. Figure 1 B and Supp. Table 1B**. The 18S was split into 4 segments before folding **Supp. Figure 1 D**, the sensitivity and specificity (the number of true positives out of total in crystal structure and the number of true positives out of the total in predicted structure) was calculated for each folded RNA. The 18S domains fold with a range of sensitivity **Suppl. Figure 1 D**. The 3’domain has the highest accuracy when folding and contains >90% of the Watson-Crick base pairs identified in the crystallography structure. 5’however has a structure with only 25% of interactions discovered. In each domain the predictions have a higher sensitivity when compared to using RNAFold alone (5’-13%, C - 19%, 3’M - 18%, 3’m **-** 79%) **Supp.Table 1C**. The PCA in **Supp. Figure 1E** shows the different structures in the ensemble for the three samples the structure highlighted with a grey box is plotted in **Supp. Figure 1F**). A subset version of this dataset is supplied with the package and the commands used in this analysis are available within the vignette of the package. The full dataset is available on GEO (GSE246412) and other COMRADES datasets can be found in previous publications (Ziv *et al*., 2018, 2020).

## 3 Conclusion

The comradesOO R package performs the analysis of data from RNA crosslinking experiments from raw reads to predicted structures. It is easy to install and portable to Windows, macOS and Linux machines with minimal installation procedure and open source. Current methods for analysing crosslinking data require performing multiple disjointed steps and manual analysis, this package solves this problem by centring around a new class, the comradesDataSet. This allows for the different data types to be stored at each stage in the analysis. We hope providing a framework for the analysis of this data will ensure that RNA crosslinking experiments are more accessible and widely adopted.

## References

Aw, J.G.A., Shen, Y., Wilm, A., Sun, M., Lim, X.N., Boon, K.-L., Tapsin, S., Chan, Y.-S., Tan, C.-P., Sim, A.Y.L., Zhang, T., Susanto, T.T., Fu, Z., Nagarajan, N. and Wan, Y. (2016) ‘In Vivo Mapping of Eukaryotic RNA Interactomes Reveals Principles of Higher-Order Organization and Regulation’, Molecular Cell, 62(4), pp. 603–617. Available at: 10.1016/j.molcel.2016.04.028.

Ban, N., Nissen, P., Hansen, J., Moore, P.B. and Steitz, T.A. (2000) ‘The complete atomic structure of the large ribosomal subunit at 2.4 A resolution’, Science (New York, N.Y.), 289(5481), pp. 905–920. Available at: 10.1126/science.289.5481.905.

Csárdi, G., Nepusz, T., Müller, K., Horvát, S., Traag, V., Zanini, F. and Noom, D. (2023) ‘igraph for R: R interface of the igraph library for graph theory and network analysis’. Zenodo. Available at: 10.5281/ZENODO.7682609.

Di Tommaso, P., Chatzou, M., Floden, E.W., Barja, P.P., Palumbo, E. and Notredame, C. (2017) ‘Nextflow enables reproducible computational workflows’, Nature Biotechnology, 35(4), pp. 316–319. Available at: 10.1038/nbt.3820.

Flynn, R.A., Zhang, Q.C., Spitale, R.C., Lee, B., Mumbach, M.R. and Chang, H.Y. (2016) ‘Transcriptome-wide interrogation of RNA secondary structure in living cells with icSHAPE’, Nature Protocols, 11(2), pp. 273–290. Available at: 10.1038/nprot.2016.011.

Fukuchi, M. and Tsuda, M. (2010) ‘Involvement of the 3’-untranslated region of the brain-derived neurotrophic factor gene in activity-dependent mRNA stabilization’, Journal of Neurochemistry, 115(5), pp. 1222–1233. Available at: 10.1111/j.1471-4159.2010.07016.x.

Kar, A., Fushimi, K., Zhou, X., Ray, P., Shi, C., Chen, X., Liu, Z., Chen, S. and Wu, J.Y. (2011) ‘RNA Helicase p68 (DDX5) Regulates tau Exon 10 Splicing by Modulating a Stem-Loop Structure at the 5′ Splice Site’, Molecular and Cellular Biology, 31(9), pp. 1812–1821. Available at: 10.1128/MCB.01149-10.

Lorenz, R., Bernhart, S.H., Höner zu Siederdissen, C., Tafer, H., Flamm, C., Stadler, P.F. and Hofacker, I.L. (2011) ‘ViennaRNA Package 2.0’, Algorithms for Molecular Biology, 6(1), p. 26. Available at: 10.1186/1748-7188-6-26.

Lu, Z., Gong, J. and Zhang, Q.C. (2018) ‘PARIS: Psoralen Analysis of RNA Interactions and Structures with High Throughput and Resolution’, Methods in molecular biology (Clifton, N.J.), 1649, pp. 59–84. Available at: 10.1007/978-1-4939-7213-5_4.

Mortimer, S.A., Kidwell, M.A. and Doudna, J.A. (2014) ‘Insights into RNA structure and function from genome-wide studies’, Nature Reviews. Genetics, 15(7), pp. 469–479. Available at: 10.1038/nrg3681.

Ray, P.S., Jia, J., Yao, P., Majumder, M., Hatzoglou, M. and Fox, P.L. (2009) ‘A stress-responsive RNA switch regulates VEGFA expression’, Nature, 457(7231), pp. 915–919. Available at: 10.1038/nature07598.

Sharma, E., Sterne-Weiler, T., O’Hanlon, D. and Blencowe, B.J. (2016) ‘Global Mapping of Human RNA-RNA Interactions’, Molecular Cell, 62(4), pp. 618–626. Available at: 10.1016/j.molcel.2016.04.030.

Shin, H., Shi, Y., Dai, C., Tjong, H., Gong, K., Alber, F. and Zhou, X.J. (2016) ‘TopDom: an efficient and deterministic method for identifying topological domains in genomes’, Nucleic Acids Research, 44(7), pp. e70–e70. Available at: 10.1093/nar/gkv1505.

Solayman, M., Litfin, T., Singh, J., Paliwal, K., Zhou, Y. and Zhan, J. (2022) ‘Probing RNA structures and functions by solvent accessibility: an overview from experimental and computational perspectives’, Briefings in Bioinformatics, 23(3), p. bbac112. Available at: 10.1093/bib/bbac112.

Spitale, R.C. and Incarnato, D. (2023) ‘Probing the dynamic RNA structurome and its functions’, Nature Reviews Genetics, 24(3), pp. 178–196. Available at: 10.1038/s41576-022-00546-w.

Tsai, M.-C., Manor, O., Wan, Y., Mosammaparast, N., Wang, J.K., Lan, F., Shi, Y., Segal, E. and Chang, H.Y. (2010) ‘Long Noncoding RNA as Modular Scaffold of Histone Modification Complexes’, Science, 329(5992), pp. 689–693. Available at: 10.1126/science.1192002.

Vicens, Q. and Kieft, J.S. (2022) ‘Thoughts on how to think (and talk) about RNA structure’, Proceedings of the National Academy of Sciences of the United States of America, 119(17). Available at: 10.1073/PNAS.2112677119/SUPPL_FILE/PNAS.2112677119.SAPP.PDF.

Wan, Y., Kertesz, M., Spitale, R.C., Segal, E. and Chang, H.Y. (2011) ‘Understanding the transcriptome through RNA structure’, Nature Reviews Genetics 2011 12:9, 12(9), pp. 641–655. Available at: 10.1038/nrg3049.

Ziv, O., Gabryelska, M.M., Lun, A.T.L., Gebert, L.F.R., Sheu-Gruttadauria, J., Meredith, L.W., Liu, Z.Y., Kwok, C.K., Qin, C.F., MacRae, I.J., Goodfellow, I., Marioni, J.C., Kudla, G. and Miska, E.A. (2018) ‘COMRADES determines in vivo RNA structures and interactions’, Nature Methods, 15(10), pp. 785–788. Available at: 10.1038/s41592-018-0121-0.

Ziv, O., Price, J., Shalamova, L., Kamenova, T., Goodfellow, I., Weber, F. and Miska, E.A. (2020) ‘The Short- and Long-Range RNA-RNA Interactome of SARS-CoV-2’, Molecular Cell, 80(6), pp. 1067–1077.e5. Available at: 10.1016/j.molcel.2020.11.004.

