## Supplemental Material for "comradesOO: An Object-Oriented R Package for the Analysis of RNA Structural Data Generated by RNA crosslinking experiments"

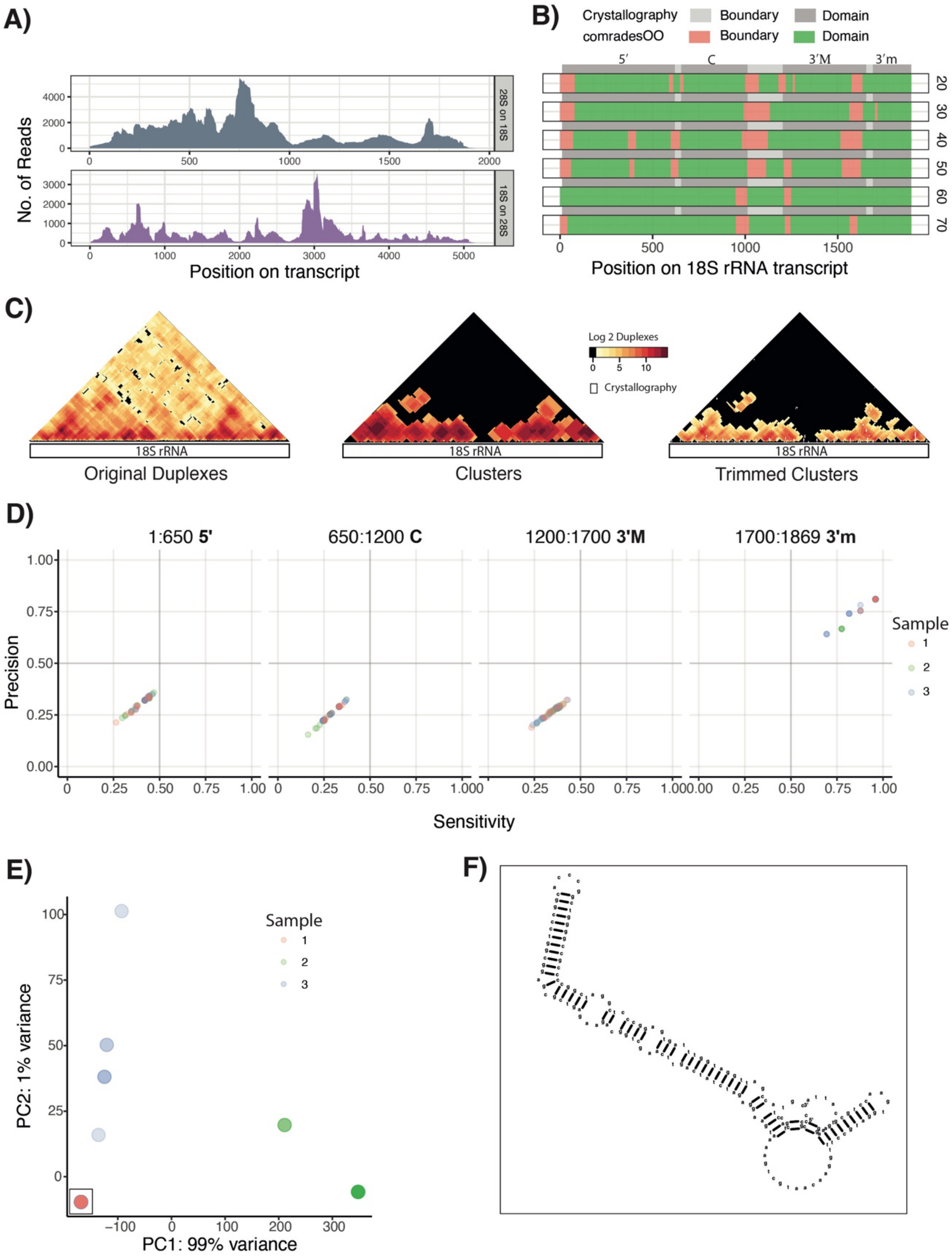


*Supplemental Figure 1.* ***Example of comradesOO analysis for the Human 18S rRNA, comparisos made with crystallography structure*** (Ban *et al.*, 2000). **A)** Intra-RNA interactions between the 28S rRNA and the 18S rRNA. **B)** Domain identification for the 18S rRNA. Grey background boxes show the domains identified in the crystallography structure **C)** Contact map showing inter-RNA interactions for the 18S RNA, raw data, clusters, and trimmed clusters. White dots on the trimmed clusters contact map shows base pairs identified by crystallography. **D)** The precision and sensitivity of each samples predicted ensemble compared to the contacts in the crystallography structure. **E)** The output from *plotEnsemlbPCA,* black box highlights the structure drawn in F. ***F)*** The output from the *plotStructure* function.

*Supplementary table 1A*. **Cluster Numbers of 18S rRNA.**

| group | sample | sampleName | rna | original | superClusters | trimmedClusters | novel | known | % known |
|---|---|---|---|---|---|---|---|---|---|
| s | 1 | s1 | RN18S1_rRNA | 125 | 83 | 83 | 18 | 65 | 0.78313253 |
| c | 1 | c1 | RN18S1_rRNA | 68 | 62 | 62 | 16 | 46 | 0.74193548 |
| s | 2 | s2 | RN18S1_rRNA | 144 | 79 | 79 | 18 | 61 | 0.7721519 |
| c | 2 | c2 | RN18S1_rRNA | 63 | 59 | 59 | 13 | 46 | 0.77966102 |
| s | 3 | s3 | RN18S1_rRNA | 133 | 85 | 85 | 17 | 68 | 0.8 |
| c | 3 | c3 | RN18S1_rRNA | 74 | 66 | 66 | 15 | 51 | 0.77272727 |

*Supplementary Table 1B.* **Output of plotDomains for 18S RRNA.**

| from.id | from.coord | to.id | to.coord | tag | size | sample | window |
|---|---|---|---|---|---|---|---|
| 176 | 1750 | 341 | 3410 | domain | 1660 | c1 | 60 |
| 342 | 3410 | 461 | 4610 | domain | 1200 | c1 | 60 |
| 462 | 4610 | 599 | 5990 | domain | 1380 | c1 | 60 |
| 600 | 5990 | 791 | 7910 | domain | 1920 | c1 | 60 |
| 792 | 7910 | 949 | 9490 | domain | 1580 | c1 | 60 |
| 950 | 9490 | 1011 | 10110 | boundary | 620 | c1 | 60 |
| 1012 | 10110 | 1210 | 12100 | domain | 1990 | c1 | 60 |
| 1211 | 12100 | 1248 | 12480 | boundary | 380 | c1 | 60 |
| 1249 | 12480 | 1615 | 16150 | domain | 3670 | c1 | 60 |
| 1616 | 16150 | 1709 | 17090 | domain | 940 | c1 | 60 |
| 1710 | 17090 | 1849 | 18490 | domain | 1400 | c1 | 60 |
| 1850 | 18490 | 1898 | 18980 | gap | 490 | c1 | 60 |
| 1 | 0 | 60 | 600 | boundary | 600 | c1 | 50 |
| 61 | 600 | 177 | 1770 | domain | 1170 | c1 | 50 |
| 178 | 1770 | 224 | 2240 | domain | 470 | c1 | 50 |
| 225 | 2240 | 376 | 3760 | domain | 1520 | c1 | 50 |
| 377 | 3760 | 401 | 4010 | boundary | 250 | c1 | 50 |
| 402 | 4010 | 456 | 4560 | domain | 550 | c1 | 50 |
| 457 | 4560 | 598 | 5980 | domain | 1420 | c1 | 50 |
| 599 | 5980 | 640 | 6400 | boundary | 420 | c1 | 50 |
| 641 | 6400 | 791 | 7910 | domain | 1510 | c1 | 50 |
| 792 | 7910 | 1011 | 10110 | domain | 2200 | c1 | 50 |
| 1012 | 10110 | 1113 | 11130 | boundary | 1020 | c1 | 50 |
| 1114 | 11130 | 1209 | 12090 | domain | 960 | c1 | 50 |
| 1210 | 12090 | 1250 | 12500 | boundary | 410 | c1 | 50 |
| 1251 | 12500 | 1354 | 13540 | domain | 1040 | c1 | 50 |
| 1355 | 13540 | 1523 | 15230 | domain | 1690 | c1 | 50 |
| 1524 | 15230 | 1583 | 15830 | boundary | 600 | c1 | 50 |
| 1584 | 15830 | 1625 | 16250 | boundary | 420 | c1 | 50 |
| 1626 | 16250 | 1713 | 17130 | domain | 880 | c1 | 50 |
| 1714 | 17130 | 1849 | 18490 | domain | 1360 | c1 | 50 |
| 1850 | 18490 | 1898 | 18980 | gap | 490 | c1 | 50 |

*Supplementary Table 1C.* **Precision and sensitivity for folding of 18S rRNA domains.**

| Shared | Total prediction | total in crytal | FP | FN | TP | Specificity | Sensitivity | START |
|---|---|---|---|---|---|---|---|---|
| 102 | 609 | 454 | 507 | 352 | 102 | 0.22 | 0.1675 | 1 |
| 27 | 205 | 142 | 178 | 115 | 27 | 0.19 | 0.1317 | 1 |
| 34 | 178 | 122 | 144 | 88 | 34 | 0.28 | 0.191 | 650 |
| 29 | 154 | 125 | 125 | 96 | 29 | 0.23 | 0.1883 | 1200 |
| 47 | 59 | 49 | 12 | 2 | 47 | 0.96 | 0.7966 | 1700 |

*Supplementary Table 2.* **Parameters and programs for read pre-processing.**

| Program | Step | Parameters |
|---|---|---|
| cuatadapt 1 | 1 | -b AGATCGGAAGAGCACACGTCTGAACTCCAGTC, -B AGATCGGAAGAGCGTCGTGTAGGGAAAGAGTGT |
| Pear | 2 | min-assembly-length 20, -v 7, and -j 5 |
| tstk collapse | 3 | --minreads 1 |
| cuatadapt 2 | 4 | -m 10, -u 6 |
| star | 5 | runThreadN 5, --genomeLoad NoSharedMemory, --outReadsUnmapped Fastx, --outFilterMultimapNmax 10000, --outFilterScoreMinOverLread 0, --outFilterMatchNminOverLread 0, --outSAMattributes All, --outSAMtype BAM SortedByCoordinate, --alignIntronMin 5, --alignIntronMax 3900000000, --seedSearchStartLmax 14, --seedSplitMin 14, --scoreGap 0, --scoreGapNoncan 0, --scoreGapGCAG 0, --scoreGapATAC 0, --scoreGenomicLengthLog2scale -0.1, --chimFilter None, --chimOutType WithinBAM HardClip, --chimSegmentMin 7, --chimJunctionOverhangMin 8, --chimScoreJunctionNonGTAG 0, --chimScoreDropMax 150, and --chimMainSegmentMultNmax 10000 |

*Supplementary Table 3.* **Column specifications for input files**

| Column Number | Description | Notes |
|---|---|---|
| 1 | Read Name | - |
| 2 | Read Sequence | - |
| 3 | Side 1 transcript ID | Works best if _ delimited and the last segment is the RNA species |
| 4 | Side 1 Position start in read sequence | - |
| 5 | Side 1 Position end in read sequence | - |
| 6 | Side 1 Coordinate start in transcript | - |
| 7 | Side 1 Coordinate end in transcript | - |
| 8 | NA | This should be "." for all rows |
| 9 | Side 2 transcript ID | Works best if _ delimited and the last segment is the RNA species |
| 10 | Side 2 Position start in read sequence | - |
| 11 | Side 2 Position end in read sequence | - |
| 12 | Side 2 Coordinate start in transcript | - |
| 13 | Side 2 Coordinate end in transcript | - |
| 14 | NA | This should be "." for all rows |
